# Peptide screening enables optimised biofunctional hydrogels for cultivated meat tissue engineering

**DOI:** 10.64898/2026.05.09.724015

**Authors:** Lea Melzener, Sergio Spaans, Christoph S. Börlin, Nicolas Hauck, Mark J. Post, Arın Doğan, Joshua E. Flack

## Abstract

Cultivated meat is an emerging biotechnology that aims to produce edible tissues in an ethical and sustainable manner. However, the recreation of skeletal muscle tissue that replicates the protein composition and sensory characteristics of traditional meat is a major challenge. Skeletal muscle tissue engineering requires non-animal-based scaffolds which are inexpensive and food-safe, while meeting specific mechanical requirements with respect to viscosity, stress-relaxation and stiffness. While many of these characteristics can be fulfilled by alginate-based biomaterials, a key limitation of alginate is its lack of intrinsic attachment sites for animal cells, preventing efficient adhesion, differentiation and tissue formation. Here, we established a screening platform to evaluate extracellular matrix (ECM)-mimicking peptides as functionalisations of alginate scaffolds in 2D. Our platform enables high-throughput assessment of cell/peptide interactions, serving as a predictive tool for 3D tissue constructs. Our screen identified two RGD-containing sequences (vitronectin- and fibronectin-mimicking peptides) as most effective in promoting attachment and myogenic fusion of bovine satellite cells. Notably, these peptides outperformed more complex mixtures containing up to seven different ECM-mimicking peptides. Our findings provide a streamlined approach for optimising biomaterial functionalisations for cultivated meat applications, and lay the groundwork for future advancements in scalable, sustainable skeletal muscle tissue engineering.

## Introduction

Cultivated meat (also referred to as ‘cultured’ meat) is an innovative biotechnology that aims to provide an alternative to traditional meat via the generation of mature muscle and fat tissues in vitro^1,2^. Interest in this technology is driven primarily by negative consequences associated with conventional meat, including greenhouse gas emissions and animal welfare concerns^3^. However, numerous scientific challenges remain to be solved before mass commercialisation can be achieved, including cost reduction, upscaling and accurate mimicry of traditional meat^2,4,5^.

The proliferation and differentiation of myogenic cells, such as muscle satellite cells (SCs) in vivo relies on close interaction between these cells and the extracellular matrix (ECM). A critical step in myogenic differentiation is the fusion of activated SCs (also referred to as myoblasts) into multinucleated myotubes, a process influenced by numerous physical and mechanical cues that affect interactions between the cell and its surroundings. These cues contribute to the generation of contractile forces which progressively generate tension within the engineered tissue, promoting muscle protein accumulation and tissue maturation^6,7^. Recreating these conditions in vitro generally necessitates the use of biomaterials (such as scaffolds and hydrogels) that can mimic these native cell/ECM interactions^8^.

Alginate is a widely used polysaccharide in tissue engineering, offering numerous advantages as a biomaterial for cultivated meat applications including its animal-free, non-toxic, and edible nature, as well as low cost and relative abundance^9–11^. Through the use of different molecular weight alginates and/or guluronic/mannuronic acid ratios, it is also possible to tune the mechanical properties of hydrogels, which can be beneficial when optimising for cell type-specific requirements^12^. ‘Fast-gelating’ hydrogel formation, in which rapid crosslinking is triggered by calcium cations (a non-toxic and food-grade method), also offers technical benefits for scalable creation of 3D constructs of various shapes and sizes, including rings or extruded fibers^13,14^. Unfortunately, as an algal-derived material, animal cells typically show little to no adhesion to native alginate, due to a lack of binding motifs. However, this lack of attachment sites can be compensated by functionalisation of the alginate backbone with specific peptide sequences^15^.

In the biomedical field, the most well-studied peptide for biomaterial functionalisation is Arg-Gly-Asp (RGD). Its wide distribution and high abundance in mammalian ECM proteins, such as collagen, and its ability to bind multiple adhesion receptors makes it suitable for many general tissue engineering applications^16,17^. However, to maximise the attachment and migration of myogenic precursors to achieve high fusion indices in alginate-based 3D tissues, cell type-specific optimisations are often necessary^8,18^.

In this study, we therefore aimed to systematically evaluate peptides from a variety of ECM proteins as functionalisations of alginate-based biomaterials, to enhance bovine satellite cell (SC) attachment and differentiation. To achieve this, we established a high-throughput 2D screening platform, allowing for the assessment of a wide range of peptides, individually and in combinations (to study potential interactions within complex mixtures) using a Design of Experiment (DoE) approach. Our simplified 2D screening platform was designed to identify promising conditions for further investigation in the context of 3D tissue environments relevant for cultivated meat production.

## Materials and Methods

### Satellite cell isolation

Muscle-derived stem cells were isolated from bovine semimembranosus muscle immediately post-slaughter, using a modified protocol previously described^19^. Briefly, muscle fibres were digested with collagenase (CLSAFA, Worthington; 1 h, 37 °C), followed by 100 μm and 40 μm cell filtration steps. Red blood cells were lysed with an Ammonium-Chloride-Potassium (ACK) lysis buffer (1 min, room temperature (RT)). Cells were plated in serum-free growth medium (SFGM, Supplementary Table 1) on fibronectin-coated (4 μg cm^-2^ bovine fibronectin; F1141, Sigma-Aldrich) tissue cultureware.

Satellite cells (SCs) were purified 72 h post-isolation using FACS as previously described^19^. Briefly, cells were stained with ITGA7-APC and ITGA5-PE and sorted using a MACSQuant Tyto Cell Sorter (Miltenyi Biotec). Flow cytometry was performed regularly during SC proliferation using a MACSQuant10 Flow Analyzer (Miltenyi Biotec) to confirm cell purity.

### Peptide coating of 2D surfaces

Pierce™ Maleimide Activated Plates (15153, Thermo Scientific) were coated with peptides following an adjusted protocol based on instructions provided by the manufacturer. Briefly, each peptide was dissolved in PBS at a 10 µM working concentration. Peptide solutions were mixed for each condition (Supplementary Tables 2, 3) in respective DoE-informed ratios. Plates were washed, incubated with peptide mixes (RT, 2 h), washed and subsequently incubated with a 10 μg ml^-1^ cysteine solution. After a final wash, SFGM was added (Supplementary Table 1). For differentiation, medium was switched to SFDM (Supplementary Table 1) after 24 h. Cells were fixed and stained for further analysis at indicated time points.

### Cell culture

SCs were proliferated in SFGM (Supplementary Table 1) on laminin-521 (0.5 μg cm^-2^, LN521-05, Biolamina) coated tissue culture vessels. After ∼12 population doublings, SCs were differentiated on peptide-functionalised tissue culture plates or on 0.5% Matrigel-coated cell culture vessels at a seeding density of 3.5×10^4^ cm^-2^. SCs were seeded in SFGM for 24 h, after which medium was exchanged to SFDM to induce myogenic differentiation (Supplementary Table 1).

### Immunofluorescent staining

Cells were fixed with 4% PFA, permeabilised with 0.5% Triton X-100 and blocked in 5% bovine serum albumin (BSA). Fixed cells were stained with rabbit α-desmin and respective secondary antibody (Supplementary Table 4) and Hoechst 33342 (Thermo Fisher Scientific), and imaged using an ImageXpress Pico Automated Cell Imaging System (Molecular Devices). Desmin area (proportion of total area that is desmin-stained) was quantified using a customised protocol within the ImageXpress Pico analysis software.

### Maleimide functionalisation of sodium alginate

1 g sodium alginate (ULV-L3, KIMICA, Japan) was dissolved in 100 mL DI water under stirring. 2.56 g (4-(4,6-dimethoxy-1,3,5-triazin-2-yl)-4-methyl-morpholinium chloride) (DMTMM, TCI, 9.25 mmol, 2 eq.) and 19.23 mg *N*-(2-aminoethyl)maleimide hydrochloride (NAMal HCl, TCI, 0.11 mmol, 0.024 eq.) were added and the reaction mixture was stirred (RT, 24 h). For purification, reaction mixture was transferred into dialysis tubing (Repligen, Spectra/Por 3 Dialysis Tubing, 3.5 kD MWCO) and dialysed against NaCl solutions with decreasing salt concentrations over the course of 24 h. During this period, dialysate was exchanged 5 times. Purified product was freeze-dried and characterised by 1H-NMR spectroscopy, showing a maleimide signal at 7.88 ppm. Quantification of maleimide functionalisation degree was performed via a modified Ellman’s test in which maleimide moieties coupled to sodium alginate were reacted with L-cysteine hydrochloride monohydrate in 0.1 M sodium phosphate buffer at pH 8.0. Excess unreacted L-cysteine hydrochloride monohydrate was photometrically quantified after reacting with Ellman’s reagent, from which the maleimide functionalisation degree of sodium alginate was 1.73 and 1.78 mol% related to alginate repeating units.

### Peptide coupling to maleimide-functionalised sodium alginate

A variety of peptides were coupled to maleimide-functionalised alginate. Peptides were synthesised by Biomatik. For each peptide to be coupled, 200 mg of maleimide-functionalised sodium alginate were dissolved in PBS at pH 7.4. All peptides were modified with a GCRD sequence at the N-terminus to react with maleimide-modified sodium alginate via thiol-Michael addition. Reaction solution was stirred (RT, 2 h), after which quantitative coupling of the peptide was assumed. For purification, reaction mixture was filled into a dialysis tubing (Repligen, Spectra/Por 3 Dialysis Tubing, 3.5 kD MWCO) and dialysed against NaCl solutions with decreasing salt concentrations for 24 h. During this period, the dialysate was exchanged 5 times. Purified alginate-peptide conjugates were freeze-dried and characterised by 1H-NMR spectroscopy showing successful conjugation of the peptide sequences.

### 3D differentiation

Peptide-functionalised alginate was dissolved to 1 wt% and mixed in respective indicated ratios (according to Fig. 4a). Peptide mixes were then mixed in a 1:1 ratio with SC suspension (5×10^7^ ml^-1^) in SFDM (Supplementary Table 1). BAMs were formed using the indicated method (Fig. 4b; Fig. 5a). Briefly, cell/alginate mixes were crosslinked in 100 mM CaCl_2_ solution for 5 min., subsequently washed in DMEM and incubated in SFDM. After 72 h (and 168 h) a full medium exchange was performed. BAMs were harvested for protein quantification and confocal microscopy at indicated time points.

### Tissue staining

BAMs were fixed with 4% paraformaldehyde (PFA) in a wash buffer containing 154 mM NaCl, 50 mM CaCl_2_ and 50 mM 3-(N-Morpholino)propanesulfonic acid, permeabilised with 0.5% Triton X-100, and blocked in 5% bovine serum albumin (BSA). Cells were stained for F-actin, desmin and myosin (Supplementary Table 4) and Hoechst 33342 (Thermo Scientific). Samples were imaged using a confocal microscope (TCS SP8, Leica Microsystems).

### Protein measurements

Protein was extracted using RIPA Lysis Buffer (sc-24948, Santa Cruz Biotechnology) after freezing at -20 °C and freeze-drying. Total protein was measured using BCA assay (ThermoFisher, 23235).

Total protein was detected (DM-TP01, Biotechne) and indicated muscle specific proteins (Supplementary Table 4) analysed using a Jess system (Biotechne).

### Design of Experiments (DoE)

#### Proliferation

DoE for testing the effect of peptide mixtures on attachment and proliferation was created as an optimal mixture design for the seven different peptides where each peptide can contribute between 0% and 100% to the total mixture. The data model was chosen as Quadratic to be able to check for direct interaction of two peptides and together with five replicate points and five lack-of-fit points resulted in the tested 38 conditions using I-Optimality.

#### Differentiation

DoE for testing the effect of peptide mixtures on differentiation was created as an optimal mixture design for the five different peptides where each peptide can contribute between 0% and 100% to the total mixture. The data model was chosen as Special Quadratic to be also able to detect non-linear interactions of three peptides and together with five replicate points and five lack-of-fit points resulted in the tested 59 conditions using I-Optimality.

#### Software and analysis

Design Expert version 13 from Stat-Ease was used to create DoEs and analyse results.

### Statistical analysis

Statistical significance was assessed using Prism v9.3.1 (GraphPad). Two-way analysis of variance was performed to determine statistical significance. Adjusted P-values used throughout the figures were: * P ≤ 0.05, ** P ≤ 0.01, *** P ≤ 0.001.

## Results

### A click chemistry platform for peptide screening in 2D and 3D (Fig. 1)

In order to investigate peptides as attachment factors for cultivated meat applications, we designed a click chemistry-based screening platform for peptide functionalisation. Maleimide-coated tissue culture plates were used for peptide screening on 2D surfaces, whilst bioartificial muscles (BAMs) were formed using maleimide-functionalised alginate for 3D experiments (Fig. 1a). Facile peptide incorporation on both culture plates and alginate backbones was achieved via the click-type reaction of maleimide and thiol groups in the cysteine residue of peptides (Fig. 1b). This Michael addition is highly reactive, selective and forms a stable thioether bond. Successful functionalisation was confirmed by 1H-NMR spectroscopy, observed by reduction of a characteristic maleimide peak (around 6.8 ppm) and the appearance of peptide-derived peaks <2.5 ppm (Supplementary Fig. 1).

**Figure 1:**
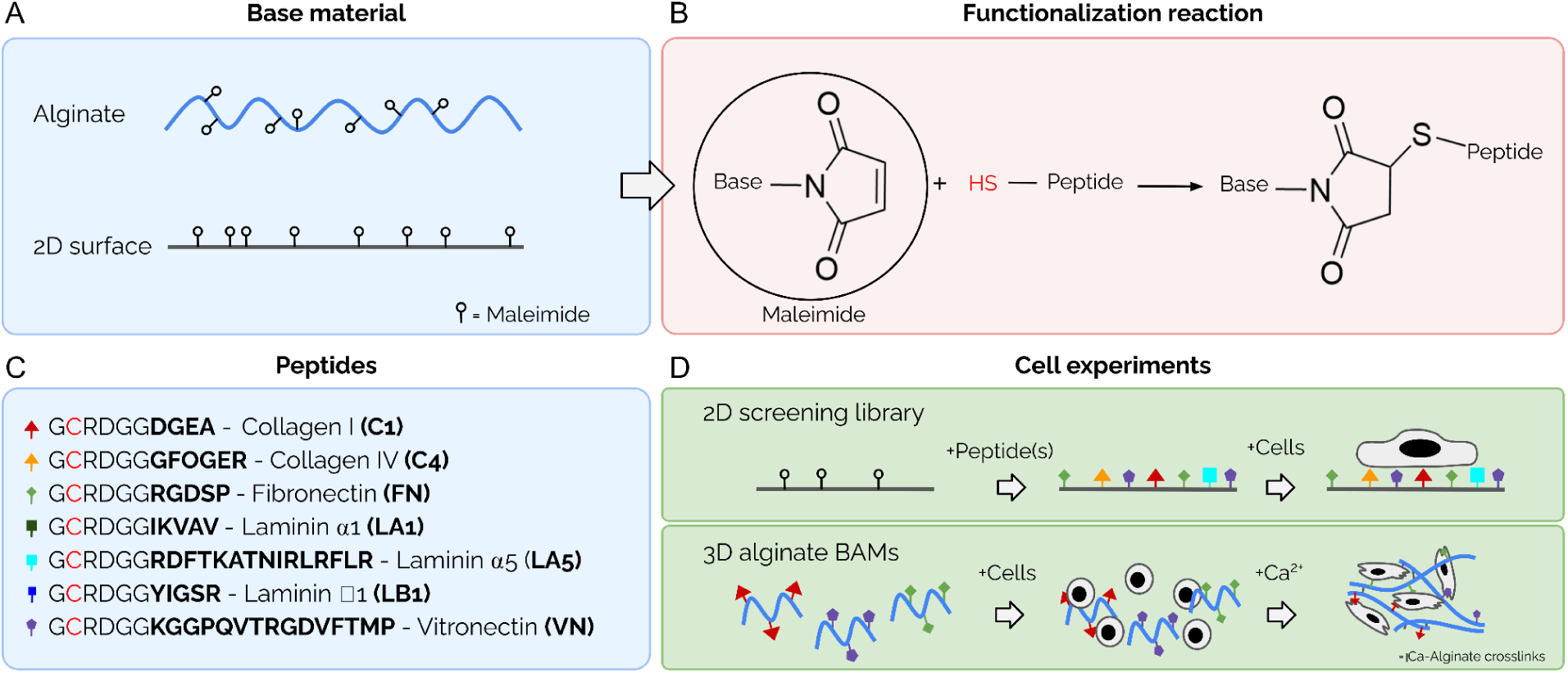
Schematic overview of click chemistry peptide functionalisation for 2D and 3D screening. A: Maleimide functionalisation of 2D surfaces and alginate backbones prior to peptide addition. B: Click-type reaction of maleimide on functionalised base material and thiol groups in cysteine residues of the peptide sequences. C: Sequences of seven selected peptides, based on reported biofunctionality and mimicry of ECM proteins. Cysteine residues required for reaction with the maleimide group are highlighted in red. D: Overview of screening approach in 2D and 3D. Alginate functionalisation with individual peptide sequences is followed by mixing in respective ratios.

Peptides were selected for inclusion in our screen based on their previously reported functionality and their mimicry of key ECM proteins, including laminin^20–22^, collagen^23,24^, fibronectin^16^ and vitronectin^25^. A total of seven peptide sequences (Fig. 1c) were incorporated into various mixtures using a Design of Experiment (DoE) framework. Peptides were systematically mixed in various ratios, and tested in cell adhesion and myogenic differentiation assays (Fig. 1d).

### VN and FN-mimicking peptides enable SC attachment in 2D (Fig. 2)

We performed a high-throughput screening experiment in 2D, in order to compare a large number of conditions with a high sensitivity for cell counts and fusion indices, and to inform subsequent experiments in the 3D system (which has a higher complexity but more closely resembles a genuine cultivated meat bioprocess).

**Figure 2:**
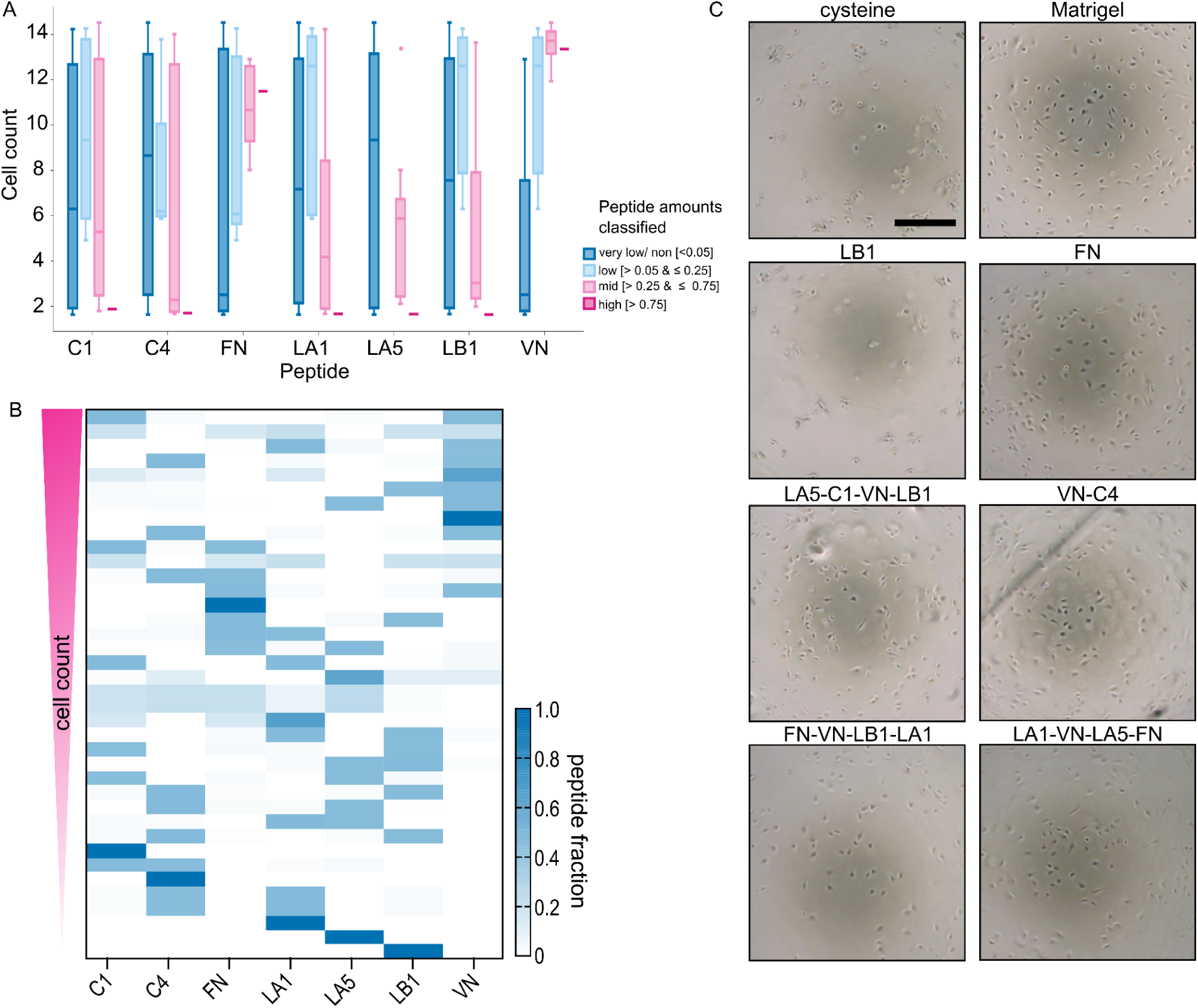
Design of Experiments (DoE)-based peptide screening for SC attachment. A: Effect of peptide amounts (as a proportion of total) on SC numbers after 72 h. B: Ranked conditions based on cell count. Shading indicates fraction of the respective peptide in the final mixture. C: Representative brightfield images of SCs on peptide-conjugated 2D surfaces 24 h after seeding. Scale bar, 200 μm.

To identify the functionality of individual peptides and potential synergistic interactions within DoE-informed mixed peptide formulations (Fig. 2a, Supplementary Table 2), bovine SC attachment and proliferation were assessed after 3 days. Peptide mixtures in which fibronectin (FN)- and vitronectin (VN)-mimicking peptides constituted the main fraction (>75%) exhibited greatest cell numbers, indicating their strong adhesive properties (Fig. 2a). In contrast, the other five peptides did not support significant cell attachment when present at similarly high concentrations (Fig. 2b). However, at lower concentrations (5 - 25%), C1, LA1, and LB1 peptides appeared to enhance cell attachment, suggesting potential synergistic effects when combined with other peptides in the mixture (Fig. 2b). Conversely, increasing the proportion of C4 and LA5 correlated with a reduction in total cell number, without showing any signs of synergistic effects with other peptide sequences (Fig. 2b), leading to their exclusion from subsequent differentiation studies.

Ranking of peptide conditions based on total cell counts at day 3 revealed that peptide mixtures with high fractions of FN-mimicking peptide and VN-mimicking peptide indeed supported greater cell attachment and proliferation (Figs. 2b, c). Both of these peptides contain RGD, suggesting that this motif promotes satellite cell adhesion. However, surfaces functionalised with VN demonstrated greater effectiveness than with FN peptide, which could be due to the context of the RGD sequence within VN (where surrounding amino acids might enhance its functionality through favourable folding or charge distribution).

### VN peptide functionalisation results in improved 2D myogenic differentiation (Fig. 3)

We subsequently assessed myogenic differentiation of SCs, again using a DoE approach based on the peptide sequences that resulted in highest cell attachment (Fig. 2). SCs were differentiated for three days, and myotube formation assessed through desmin staining^26^.

**Figure 3:**
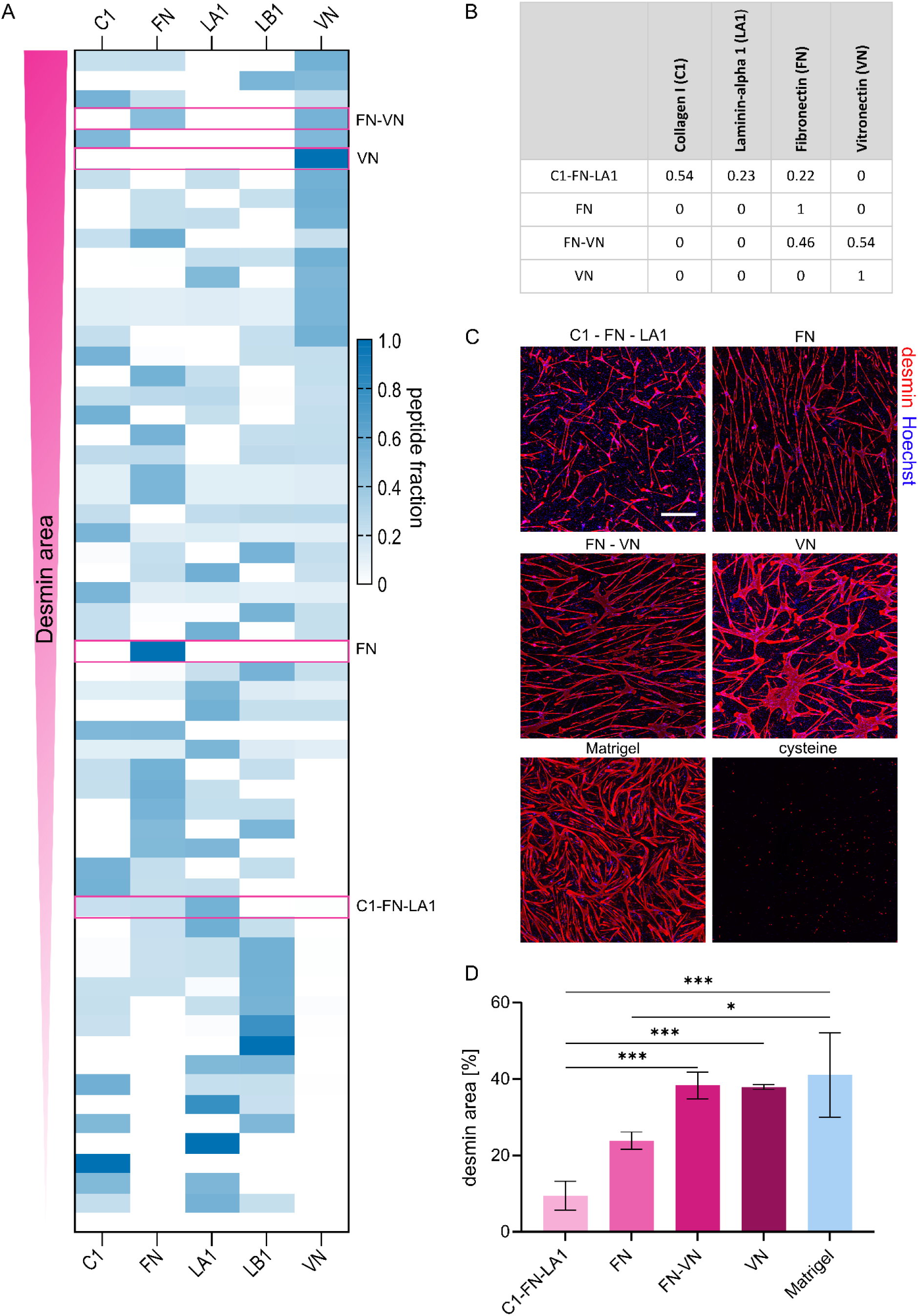
VN peptide functionalisation results in improved 2D myogenic differentiation. A: Ranked conditions based on desmin area. Colour indicates fraction of respective peptide in the final mix. B: Table of representative peptide fractions in mixes. C: Fluorescence images of differentiating SCs to visualise myogenic fusion after 72 h on indicated peptide coatings. Blue, Hoechst; red, desmin. Scale bar, 500 µm. D: Mean quantified desmin area (proportion of total image field that is desmin-stained) for images in B. Error bars indicate s.d., n = 3. *P ≤ 0.05, **P ≤ 0.01, ***P ≤ 0.001.

Ranking the conditions in terms of desmin-positive area revealed that higher fractions of VN peptide led to improved differentiation (Fig. 3a). Consistent with the attachment screening, C1, LA1, and LB1 peptides exhibited reduced myogenic fusion when present at high concentrations in the functionalisation mix. In contrast, FN peptide alone supported moderate differentiation, which was further enhanced by the addition of VN (Fig. 3a). Quantification correlated with visual observation of the desmin-stained images (Fig. 3c).

While the quantified desmin-positive area was comparable between FN-VN, VN and Matrigel control coated surfaces (Fig. 3d), myotubes showed distinct morphological differences (Fig. 3c), which could hint towards differences in binding affinities and quantity of attachment sites on these surfaces. While Matrigel-coated surfaces resulted in a homogeneous myotube morphology with a high degree of branching, peptide-functionalised surfaces showed more heterogeneous myotube formation, with reduced branching and networking between myotubes. Furthermore, morphological differences between the different peptide coatings could be observed, with FN-VN peptide coated surfaces resulting in thin myotubes, while VN peptide coated surfaces showed wider structures (Fig. 3c).

### Altering peptide functionalisation yields improved cell/alginate interactions (Fig. 4)

Following the identification of VN- and FN-mimicking peptides as the most effective sequences for promoting cell adhesion and differentiation in 2D, a subset of conditions was selected for validation in a 3D alginate hydrogel system (Fig. 3b). VN and FN peptides were tested individually and in combination, while non-functionalised alginate and a low-performing peptide mix from the 2D tests (C1-LA5-FN) served as negative controls (Fig. 3b).

Low molecular weight alginate functionalised with the selected peptides was combined with cells and crosslinked in a calcium chloride solution, resulting in torus-shaped structures. Over time, hydrogels underwent compaction as cells adhered to the biomaterial (Fig. 4a). Immunofluorescence staining on day 7 revealed elevated levels of desmin and phalloidin in the FN, VN, and FN-VN conditions. VN and FN-VN peptide functionalisation resulted in the formation of elongated myotubes, whereas C1-LA5-FN and FN conditions predominantly led to the development of skeletal muscle protein-enriched spherical structures, which we termed ‘myospheres’. These findings were consistent across cells derived from three donor animals (Fig. 4b). Non-functionalised low molecular weight alginate was excluded from subsequent analysis, as it dissolved fully within the first 4 days (due to the absence of cell/hydrogel) interactions. Conversely, all peptide-functionalised conditions maintained compacted tissue constructs, suggesting successful functionalisation resulting in various degrees of cell/hydrogel interaction.

**Figure 4:**
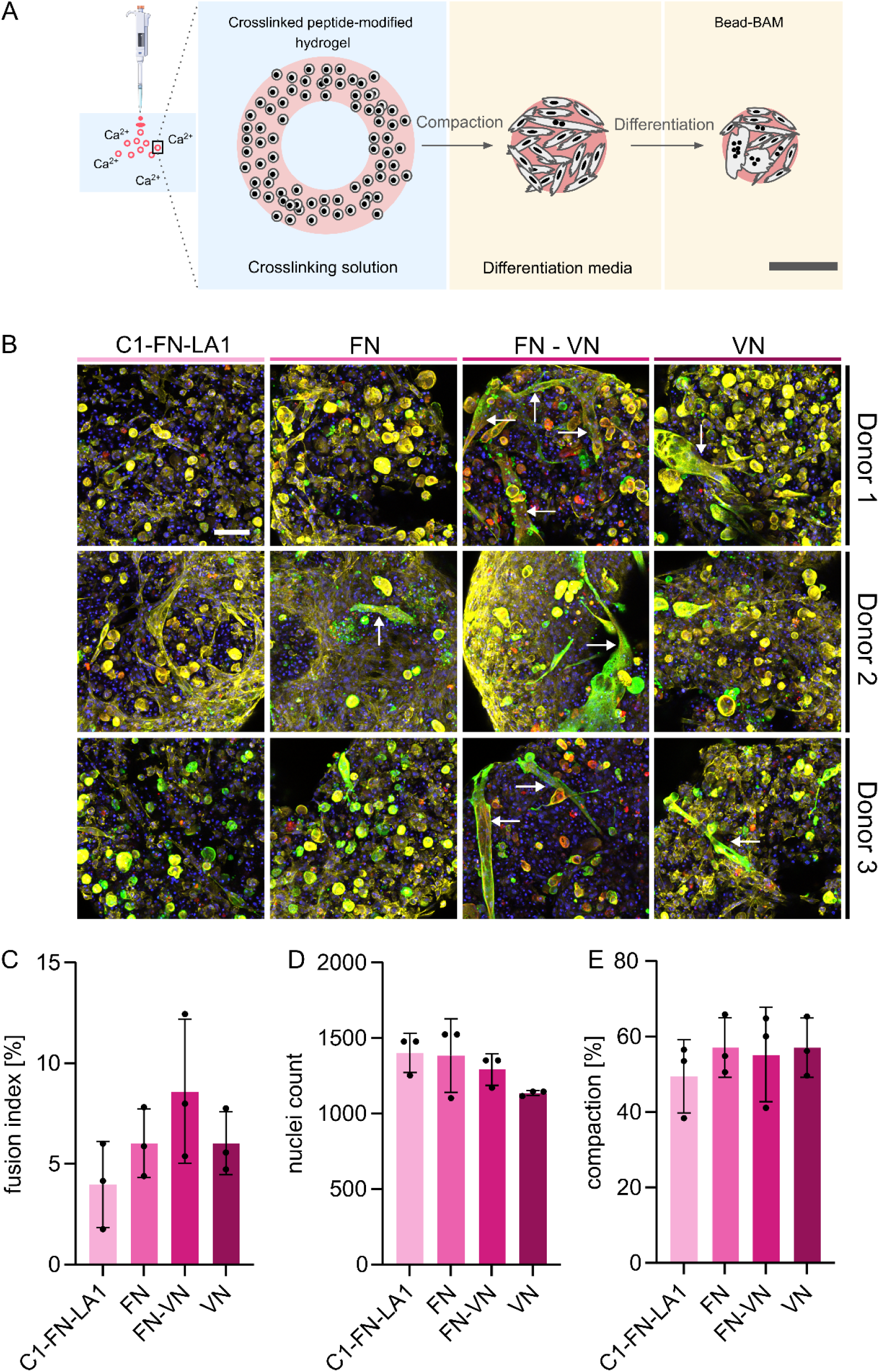
Altering peptide functionalisation yields improved cell/alginate interactions. A: Schematic representation of bead-BAM formation. B: Fluorescence images of differentiating SCs in bead-BAMs to visualise myogenic fusion after 168 h in alginate functionalised with indicated peptides. Blue, Hoechst; yellow, phalloidin; green, desmin; red, myosin. Arrows indicate elongated myotubes. Scale bar, 200 µm. C: Mean quantified myosin and desmin based fusion indices for images in B. Error bars indicate s.d., n = 3. No statistically significant differences were observed, with all P-values > 0.05. D: Mean quantified nuclei counts for images in B. Error bars indicate s.d., n = 3. No statistically significant differences were observed, with P-values > 0.05. E: Mean quantified bead-BAM compaction after 168 h. Error bars indicate s.d., n = 3. No statistically significant differences were observed, with P-values > 0.05.

The mixture of FN and VN peptides yielded the highest differentiation, as indicated by cell morphology and fusion rates, followed by pure FN peptide and VN peptide (Figs. 4b, c). Peptide mixtures containing VN peptide were associated with slightly reduced nuclear counts (Fig. 4d). The lowest fusion index was observed with C1-LA5-FN (Fig. 4c), correlating with slower compaction and suggesting reduced cell/hydrogel interactions (Fig. 4e).

2D results (Figs. 2, 3) were largely predictive for successful tissue formation and differentiation in 3D (Fig. 4). However, in 3D, the distinction between VN and FN (Figs. 3a-c) was not so evident, while their combination (FN-VN) significantly enhanced differentiation (Figs. 4b, c). VN-functionalised alginate did not result in increased fusion rates in 3D, but contributed to improved cell morphology, promoting the formation of aligned myotubes (Fig. 4b). Overall, our data suggest that lack of cell adhesion to the FN peptide contributes to the formation of ‘myosphere’ structures, in which significant further maturation is unlikely. VN, on the other hand, led to the formation of elongated myotubes, which seems more promising for tissue maturation^27,28^.

### Myotube self-anchorage does not improve maturation (Fig. 5)

Following the improvement of myogenic differentiation through vitronectin (VN) peptide functionalisation in both 2D and 3D culture systems, we aimed to investigate whether peptide modification could prolong the duration of elongated myotubes in a timecourse experiment. Additionally, since myogenic differentiation and maturation can be promoted by mechanical anchorage, we introduced an additional culture system in which the hydrogel mixtures compact around stainless steel pillars (Fig. 5a).

**Figure 5:**
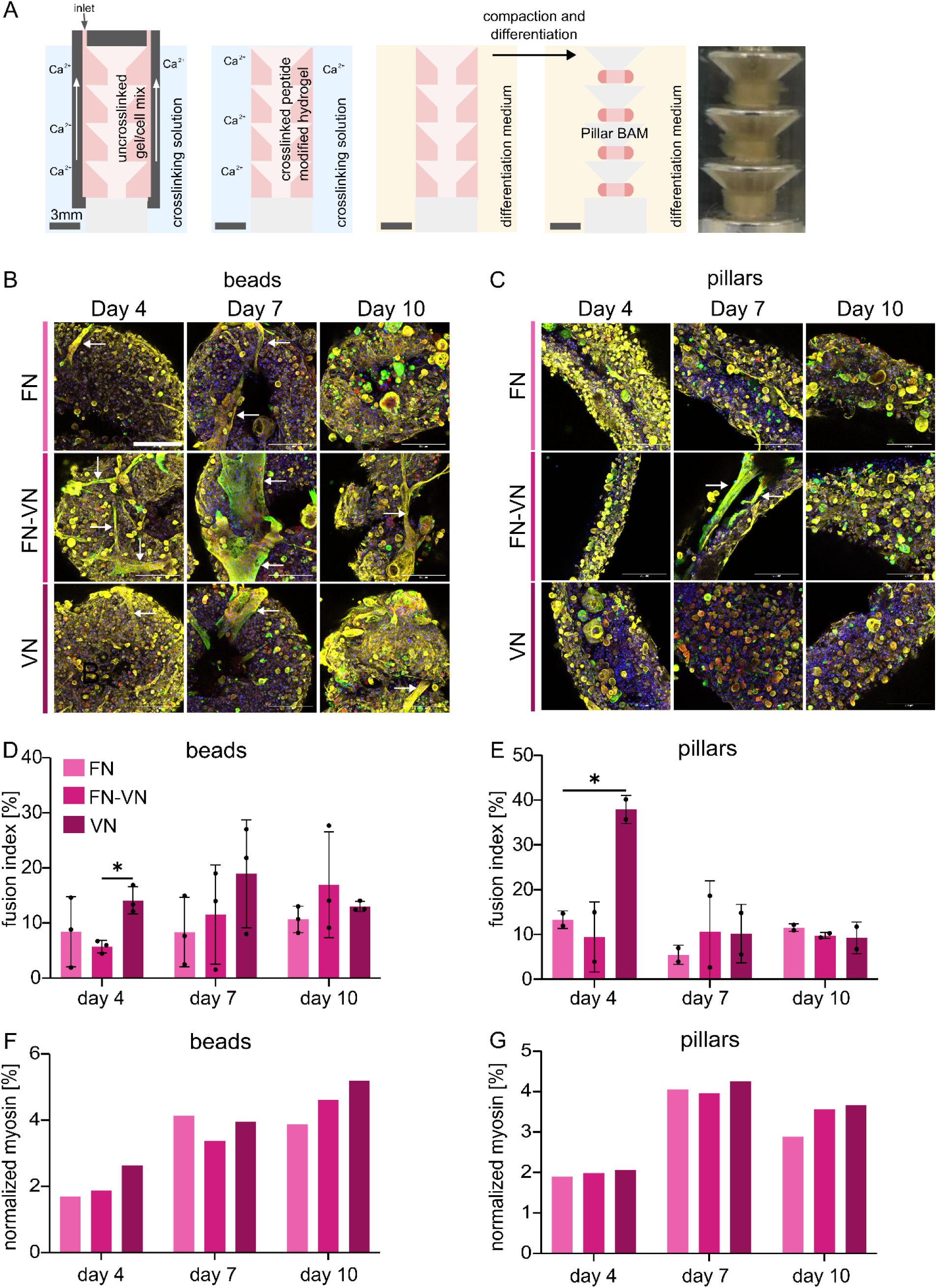
Myotube self-anchorage in pillar BAMs does not improve maturation. A: Schematic of pillar-BAM formation. B: Fluorescence images of differentiating SCs in bead-BAMs to visualise myogenic fusion at indicated timepoints in alginate functionalised with indicated peptide sequences. Blue, Hoechst; yellow, phalloidin; green, desmin; red, myosin. Arrows indicate myotubes. Scale bar, 200 µm. C: Fluorescence images of differentiating SCs in pillar-BAMs to visualise myogenic fusion at indicated timepoints in alginate functionalised with indicated peptides. Blue, Hoechst; yellow, phalloidin; green, desmin; red, myosin. Arrows indicate myotubes. Scale bar, 200 µm. D: Mean quantified myosin- and desmin-based fusion indices for images in B. Error bars indicate s.d., n = 3. E: Mean quantified myosin- and desmin-based fusion indices for images in C. Error bars indicate s.d., n = 3. F: Myosin expression normalised to a meat lysate control, analysed by western blot after 4, 7 and 10 days differentiation in indicated peptide-functionalised bead BAMs. G: As F, but in functionalised alginate pillar BAMs. *P ≤ 0.05.

Cells were mixed with three distinct peptide-functionalised alginate formulations; FN, FN-VN, and VN, and subsequently crosslinked by dripping the cell-laden mixtures into calcium chloride solution to form beads (Fig. 4b), or by moulding it around pillars (Fig. 5a). Immunofluorescence staining on days 4, 7, and 10 revealed that myotube formation occurred earlier in FN-VN beads compared to FN and VN alone (Fig. 5b). Across all three conditions, both elongated myotubes and the ‘myospheres’ were observed, with FN-VN demonstrating the highest proportion of myotubes (Fig. 5b). However, no clear trends could be observed regarding increased maturation in any of the peptide sequences or culture systems (Figs. 5b-g). Fusion indices stabilised or decreased in all peptide conditions over time in both beads and pillar BAMs (Figs. 5b-d). Despite this, myosin quantification revealed trends towards an increased expression up to day 10 in bead BAMs across all peptide conditions (Fig. 5f). In pillar BAMs, myosin expression peaked on day 7 and decreased afterwards (Fig. 5g). These data suggest that myogenic fusion occurs only during the early stages of tissue formation, but that maturation of those cells that do fuse can continue at later timepoints.

## Discussion

The generation of 3D constructs in which stem cells can differentiate to produce mature skeletal muscle tissue that mimics the appearance, texture, taste and nutritional value of traditional meat is a key challenge in cultivated meat development. However, this requires extensive optimisation of non-animal-based biomaterials to improve cellular attachment and differentiation. Here we developed a simple 2D culture platform that can be used as a tool to efficiently screen for peptide sequences that offer promise as biomaterial functionalisations. We used this system to identify several sequences that improve SC attachment and myotube formation in 3D alginate hydrogel-based bioartificial muscles (BAMs).

RGD, a well-known motif for cell attachment, can be found in a broad range of extracellular matrix proteins (including collagen and fibronectin), and is often used to functionalise biomaterials for tissue engineering purposes^16,29^. However, whilst providing cell attachment sites for SCs, we previously observed that myotubes fail to mature in RGD-alginates^26,30^. This may be due to the limited extent of initial cell attachment or changing adhesion requirements during the process of myogenic differentiation, amongst other factors. To improve maturation, we therefore explored a broader range of peptides, designed to mimic a variety of ECM proteins. By adding these to maleimide-coated plates using click-chemistry, either individually or in combination, our 2D platform allows for quick and easy assessment of cell attachment and differentiation. We were able to identify a vitronectin-mimicking sequence (VN) that resulted in higher SC attachment and fusion when compared to RGD. This may be due to the strong binding affinity of SCs to vitronectin via α_V_β_3_ integrin^31^. A more detailed study using transcriptomics or proteomics in different biomaterial contexts could reveal further patterns of SC integrin expression, which could then inform peptide design for further optimisation^19,32^. Although the laminin peptides tested in this study did not result in cell attachment, the high expression of laminin-associated integrins such as ITGA7 and ITGB1 in SCs suggests that further exploration of laminin-mimicking sequences might still be valuable^19^.

Our 2D platform proved effective for the screening of both individual peptides and combinations of up to seven peptide sequences in the context of a DoE-based experimental design. In future studies, larger arrays of rationally or semi-rationally designed sequences can be tested in an efficient fashion. The FN- and VN-mimicking peptides which showed the most promising results were the only sequences containing an RGD motif. It would therefore be interesting to incorporate RGD motifs within other sequences, and to explore further RGD-containing sequences in which flanking amino acids can add additional functionality for cell binding, including charges^33^ and/or peptide configuration. Our VN peptide contains a positively charged lysine, which has previously been shown to promote SC adhesion^34^. Flanking amino acids can also affect peptide configuration, with knock-on effects on integrin binding affinity^35^. For example, cyclic-RGD (a well-known example used to improve cell adhesion compared to linear RGD) closely resembles the natural configuration of RGD found in fibronectin^36,37^.

Positive results for peptides in our 2D screening platform generally translated to favourable performance in 3D BAMs compared to RGD-alginate alone. However, considerable variability was also observed in terms of both the extent and the consistency of differentiation, suggesting that much further optimisation is still required. Our frequent observation of spherical ‘myosphere’ structures enriched in skeletal muscle proteins, rather than elongated myotubes, suggests insufficient attachment to the biomaterial that results in ‘collapse’ of myotubes as a contractile apparatus is assembled during differentiation (as has previously been observed^26,32^). Numerous additional aspects of 3D tissue culture, including biomaterial stiffness and porosity, as well as BAM ultrastructure, can also affect cell adhesion, tension generation and maturation of myotubes^38^. We investigated the effect of BAM ultrastructure by comparing an anchor-free system (beads) with a system that allows self-anchoring (pillars). The additional anchoring of myotubes had no significant effect on myotube structure or differentiation in our system, even though previous reports have suggested that anchoring promotes cell alignment, contributing to maturation of muscle tissue^39^. VN peptide functionalisation, which led to greater numbers of elongated myotubes in anchorage-free BAM systems in our experiments, thus offers a promising basis for maturation in optimised macrostructures. In the context of a cultivated meat bioprocess, striking the balance between scalable biomanufacturing and optimal tissue formation is challenging. Alternative shapes for tissue constructs, such as fibres formed via extrusion-based systems, might also be investigated to better understand the effect(s) that BAM ultrastructure plays on tissue formation.

### Conclusion

Overall, our results indicate that the production of 3D skeletal muscle constructs that genuinely mimic traditional meat can be improved through the tuning of both individual peptide sequences, and the overall composition in a peptide mixture. 2D experiments can effectively identify promising peptides for testing in 3D tissues, and our platform can thus be used to efficiently screen many further candidates. Nevertheless, significant further work remains to both understand the contribution of different ECM-mimicking peptides to SC attachment and differentiation, and to develop a full understanding of the biology that is essential for the creation of fully mature 3D tissues in vitro.

## Availability of data and materials

Data supporting findings of this study are available from the authors on request.

## Author contributions

LM, SS and NH designed and performed experiments, and analysed the data. CB designed experiments and analysed data. MJP, AD and JEF supervised the study, designed experiments and analysed data. LM and JEF wrote the manuscript with input from all authors.

## Acknowledgements

We would like to thank André Pötgens for assistance with protein measurements, Joseph Caponi for automated liquid handling, and Dirk Remmers for code and technical assistance with image analysis. We thank Rui Hueber for help with cell isolation. We would also like to acknowledge all members of the Cell Biology and Muscle Tissue Development teams of Mosa Meat for their helpful discussions during the design, execution and analysis of this study.

