## Supplementary Material File for "Peptide screening enables optimised biofunctional hydrogels for cultivated meat tissue engineering"

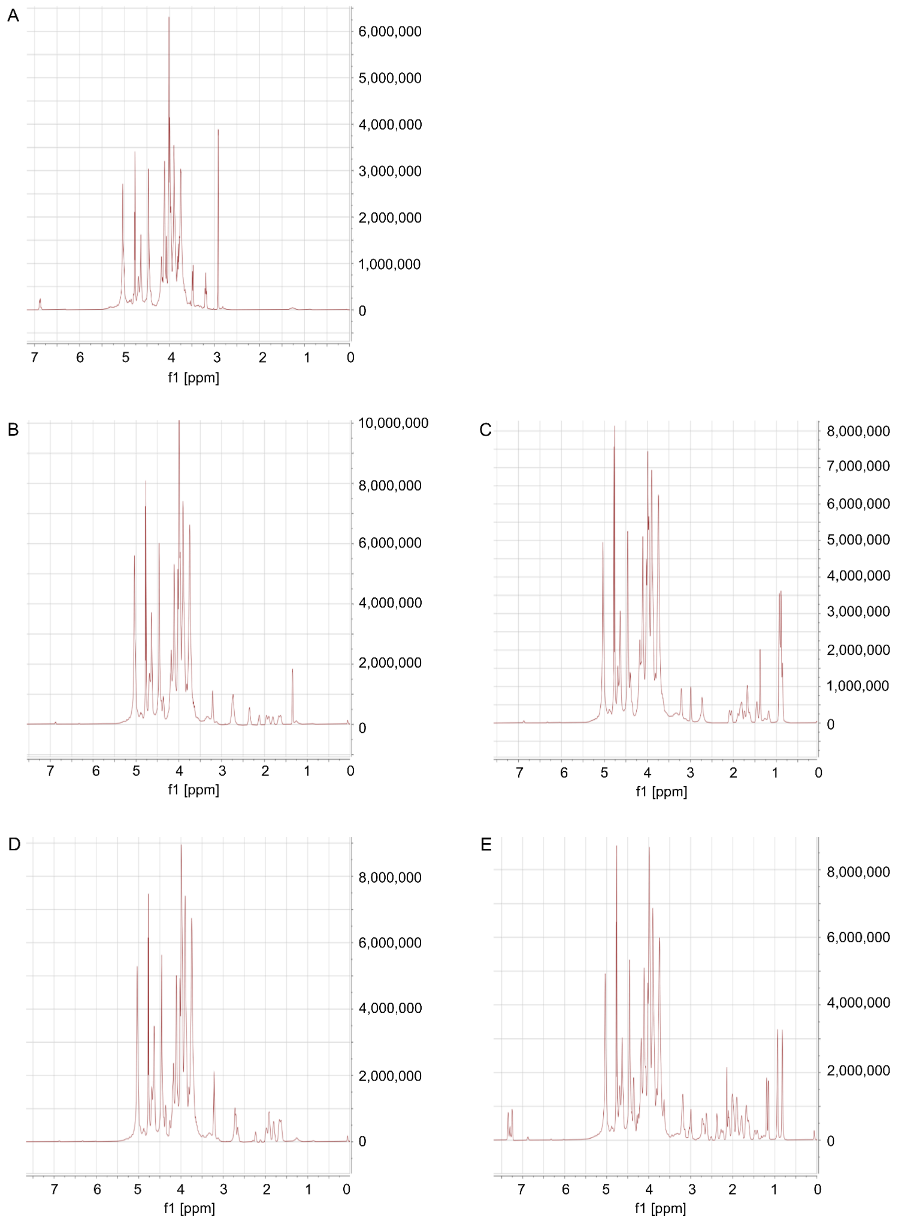


**Supplementary Figure 1:**

A: 1H-NMR spectrum of maleimide functionalization recorded in D2O; DS by Ellman's: 1.73 mol% related to sodium alginate repeating units.

B: 1H-NMR spectrum of collagen I peptide functionalization recorded in D2O; DS by Ellman's: 1.73 mol% related to sodium alginate repeating units, GCRDGGDGEA (SP210671).

C: 1H-NMR spectrum of laminin alpha-5 peptide functionalization recorded in D2O; DS by Ellman's: 1.73 mol% related to sodium alginate repeating units, GCRDGGRDFTKATNIRLRFLR (SP210671).

D: H-NMR spectrum of fibronectin peptide functionalization recorded in D2O; DS by Ellman's: 1.73 mol% related to sodium alginate repeating units, GCRDGGRGDSP (SP210671).

E: H-NMR spectrum of vitronectin peptide functionalization recorded in D2O; DS by Ellman's: 1.73 mol% related to sodium alginate repeating units, GCRDGGKGGPQVTRGDVFTMP (SP210671).

**Supplementary Table 1: Medium formulations**

| Compound | Supplier | Concentration |
|---|---|---|
| **Serum free growth medium (SFGM)** | | |
| DMEM/F-12 | P04-041262B, PAN Biotech | |
| α-linolenic acid | L2376, Sigma Aldrich | 1.0 μg $ml^{-1}$ |
| bFGF-2 | 100-18B, Peprotech | 10 ng $ml^{-1}$ |
| Bovine Serum Albumin (BSA) | A9418, Sigma Aldrich | 5.0 mg $ml^{-1}$ |
| bHGF | 100-39H, Peprotech | 50 ng $ml^{-1}$ |
| Hydrocortisone | H0888, Sigma Aldrich | 0.1 mM |
| Insulin, Transferrin, Selenium, Ethanolamine (ITSE) | 00-101, biogems | 1% |
| GlutaMax | 35050061, ThermoFisher | 2 mM |
| D-glucose | G7021, Sigma Aldrich | 17.7 mM |
| L-ascorbic acid 2-phosphate (Vitamin C) | A8960, Sigma Aldrich | 155 μM |
| Penicillin/Streptomycin/Amphotericin (PSA) | 17-745E, Lonza | 1% |
| T3 | T6397, Sigma Aldrich | 20 ng $ml^{-1}$ |
| **Serum free differentiation medium (SFDM)** | | |
| DMEM | A14430-01, Gibco | |
| EGF-1 | AF-100-15, Peprotech | 10 ng $ml^{-1}$ |
| D-glucose | G7021, Sigma | 5.5 mM |
| GlutaMax | 35050061, ThermoFisher | 2 mM |
| Human Serum Albumin | Rc HA NW20, Richcore Lifesciences | 0.5 mg $ml^{-1}$ |
| ITSE | 00-101, biogems | 2% |
| L-ascorbic acid 2-phosphate (Vitamin C) | A8960, Sigma Aldrich | 40 μM |
| Lysophosphatidic acid (LPA) | L7260, Sigma Aldrich | 1 μM |
| MEM Amino Acids Solution | 11130-051, ThermoFisher | 0.50% |
| $NaHCO_3$ | P2256, Sigma Aldrich | 6.5 mM |
| PSA | 17-745E, Lonza | 1% |
| Soy hydrolysates | 58903C, Merck | 1% |
| Sodium l-lactate | 71718, Sigma | 10 mM |
| Sodium pyruvate | P2256, Sigma Aldrich | 0.5 mM |

### Supplementary Table 2: Design of Experiment peptide mixes for attachment study (Fig. 2)

| GCRDGGRGDSP (SP210671) | GCRDGGDGEA (SP210671) | GCRDGGGFOGER (SP210671) | GCRDGGRGDSP (SP210671) | GCRDGGIKVAV (SP210671) | GCRDGGRDFTKATNIRLRFLR (SP210671) | GCRDGGYIGSR (SP210671) | GCRDGGKGGPQVTRGDVFTMP (SP210671) |
|---|---|---|---|---|---|---|---|
| FN | C1 | C4 | FN | LA1 | LA5 | LB1 | VN |
| 0 | 0 | 0 | 0 | 0 | 0 | 1 | 0 |
| 0 | 0 | 0.481374 | 0 | 0.029524 | 0 | 0.0203689 | 0.468733 |
| 0.162796 | 0.167825 | 0 | 0.162796 | 0.669379 | 0 | 0 | 0 |
| 0 | 0.479063 | 0.469068 | 0 | 0.0194773 | 0.0323926 | 0 | 0 |
| 0 | 0 | 0 | 0 | 1 | 0 | 0 | 0 |
| 0.484468 | 0 | 0 | 0.484468 | 0.025 | 0 | 0.0258642 | 0.464668 |
| 0.0030179 | 0.022628 | 0.0259379 | 0.0030179 | 0 | 0.466192 | 0 | 0.482224 |
| 0 | 0.0236765 | 0.0165534 | 0 | 0.488075 | 0.471695 | 0 | 0 |
| 0.00776669 | 0.025 | 0.476352 | 0.00776669 | 0 | 0.02 | 0.470881 | 0 |
| 0 | 0.00236441 | 0 | 0 | 0.0253373 | 0.471889 | 0.480492 | 0.019918 |
| 0.483386 | 0 | 0.02 | 0.483386 | 0 | 0.03 | 0.466614 | 0 |
| 0.00776669 | 0.025 | 0.476352 | 0.00776669 | 0 | 0.02 | 0.470881 | 0 |
| 0 | 1 | 0 | 0 | 0 | 0 | 0 | 0 |
| 0.0355586 | 0.463654 | 0 | 0.0355586 | 0.027146 | 0 | 0.473641 | 0 |
| 0.223514 | 0.198299 | 0.222877 | 0.223514 | 0.0762842 | 0.259026 | 0.02 | 0 |
| 0 | 0.0175282 | 0.123941 | 0 | 0 | 0.627559 | 0.113946 | 0.117026 |
| 0 | 0.476592 | 0 | 0 | 0.475391 | 0 | 0.0105452 | 0.0374719 |
| 0 | 0.02 | 0.470054 | 0 | 0.479946 | 0 | 0.03 | 0 |
| 0 | 0.481587 | 0.0258501 | 0 | 0 | 0.02 | 0 | 0.472563 |
| 0.468363 | 0.486009 | 0.0181019 | 0.468363 | 0 | 0 | 0 | 0.0275263 |
| 0.474141 | 0.0155155 | 0.476606 | 0.474141 | 0 | 0 | 0 | 0.0337375 |
| 0.159382 | 0.20366 | 0 | 0.159382 | 0.224049 | 0 | 0.203016 | 0.209892 |
| 0.480169 | 0.0285096 | 0.0245423 | 0.480169 | 0.466779 | 0 | 0 | 0 |
| 0 | 0 | 0.00104232 | 0 | 0 | 0 | 0 | 0.998958 |
| 0.025 | 0 | 0 | 0.025 | 0.477224 | 0 | 0.47364 | 0.0241362 |
| 0 | 0.02 | 0.470054 | 0 | 0.479946 | 0 | 0.03 | 0 |
| 0 | 0.130842 | 0.0728635 | 0 | 0.149768 | 0 | 0 | 0.646526 |
| 0.026812 | 0 | 0.480672 | 0.026812 | 0.02 | 0.467516 | 0.005 | 0 |
| 0 | 0.0345087 | 0.02 | 0 | 0 | 0 | 0.460371 | 0.485121 |
| 0 | 0 | 0.481374 | 0 | 0.029524 | 0 | 0.0203689 | 0.468733 |
| 0 | 0 | 0 | 0 | 0 | 1 | 0 | 0 |
| 0.00842538 | 0 | 0 | 0.00842538 | 0.482005 | 0.0489026 | 0 | 0.460667 |
| 1 | 0 | 0 | 1 | 0 | 0 | 0 | 0 |
| 0.223514 | 0.198299 | 0.222877 | 0.223514 | 0.0762842 | 0.259026 | 0.02 | 0 |
| 0.461535 | 0 | 0 | 0.461535 | 0.025 | 0.483245 | 0 | 0.0302196 |
| 0 | 0 | 1 | 0 | 0 | 0 | 0 | 0 |
| 0.025 | 0.464545 | 0 | 0.025 | 0 | 0.469638 | 0.0408175 | 0 |
| 0.159382 | 0.20366 | 0 | 0.159382 | 0.224049 | 0 | 0.203016 | 0.209892 |

**Supplementary Table 3: Design of Experiment peptide mixes for differentiation study (Fig. 3)**

| GCRDGGDGEA (SP210671) | GCRDGGRGDSP (SP210671) | GCRDGGIKVAV (SP210671) | GCRDGGYIGSR (SP210671) | GCRDGGKGGPQVTRG DVFTMP (SP210671) |
|---|---|---|---|---|
| C1 | FN | LA1 | LB1 | VN |
| 0.5536691732 | 0 | 0.2207081843 | 0.2243693435 | 0.001253299012 |
| 0.2278969309 | 0 | 0 | 0.2200859826 | 0.5520170865 |
| 0 | 0.4843891473 | 0 | 0.5156108527 | 0 |
| 0 | 0 | 0.4962587497 | 0.5037412503 | 0 |
| 0 | 0.4644655595 | 0 | 0 | 0.5355344405 |
| 0.2206549104 | 0.008453594773 | 0.01191794662 | 0.5343600528 | 0.2246134955 |
| 0.4921700043 | 0 | 0.5078299957 | 0 | 0 |
| 0.1177129242 | 0.1248073665 | 0.5103745668 | 0.1315530102 | 0.1155521324 |
| 0 | 0.2184566702 | 0.5578806613 | 0.2236626685 | 0 |
| 0.5211647968 | 0.1206602969 | 0.1336185828 | 0.1102110357 | 0.1143452877 |
| 0.2248903704 | 0 | 0.223606653 | 0.5327038974 | 0.01879907922 |
| 0 | 0.5360683082 | 0 | 0.2273969653 | 0.2365347265 |
| 0.498931018 | 0.501068982 | 0 | 0 | 0 |
| 0 | 0.492464705 | 0.507535295 | 0 | 0 |
| 1 | 0 | 0 | 0 | 0 |
| 0.2109780268 | 0 | 0.01792335393 | 0.767860561 | 0.003238058243 |
| 0.2326381524 | 0 | 0.2397601273 | 0.2606489297 | 0.2669527906 |
| 0 | 0 | 1 | 0 | 0 |
| 0 | 0 | 0 | 1 | 0 |
| 0 | 0.2247729984 | 0.2334390256 | 0 | 0.541787976 |
| 0.2186879165 | 0.2296605001 | 0.545149578 | 0 | 0.006502005412 |
| 0 | 1 | 0 | 0 | 0 |
| 0 | 0 | 0 | 0 | 1 |
| 0 | 0.234451103 | 0.5539425086 | 0 | 0.2116063884 |
| 0.5211647968 | 0.1206602969 | 0.1336185828 | 0.1102110357 | 0.1143452877 |
| 0.2365530291 | 0.2642446646 | 0.2725991726 | 0.007144072929 | 0.2194590608 |
| 0.1189382842 | 0.514371599 | 0.118404698 | 0.11908393 | 0.1292014888 |
| 0.1189382842 | 0.514371599 | 0.118404698 | 0.11908393 | 0.1292014888 |
| 0.5393321818 | 0.008095940905 | 0 | 0.2247896007 | 0.2277822766 |
| 0 | 0 | 0 | 0.517926614 | 0.482073386 |
| 0.1177129242 | 0.1248073665 | 0.5103745668 | 0.1315530102 | 0.1155521324 |
| 0.5421771324 | 0.2242000016 | 0.233622866 | 0 | 0 |
| 0.02159878886 | 0.2204530779 | 0 | 0.5305354532 | 0.22741268 |
| 0.2248132379 | 0.5538266083 | 0.2213601538 | 0 | 0 |
| 0 | 0.02342359995 | 0.2131606404 | 0.5389165936 | 0.2244991661 |
| 0.5 | 0 | 0 | 0 | 0.5 |
| 0.1253980405 | 0.1151854478 | 0.1282873892 | 0.1098044354 | 0.5213246871 |
| 0.2197405924 | 0.5648933838 | 0 | 0 | 0.2153660238 |
| 0 | 0 | 0.5522978638 | 0.2238617111 | 0.2238404251 |
| 0.5396954187 | 0.2251688667 | 0 | 0 | 0.2351357146 |

| | | | | |
|---|---|---|---|---|
| 0.01739861546 | 0.2088879599 | 0.2223130609 | 0.5471158311 | 0.004284532612 |
| 0 | 0.005892429838 | 0.225930574 | 0.2159071599 | 0.5522698362 |
| 0.228952177 | 0.5412720859 | 0 | 0.2297757371 | 0 |
| 0.2250233724 | 0 | 0.2247702606 | 0.003666583254 | 0.5465397837 |
| 0 | 0.5421829807 | 0.2263083728 | 0 | 0.2315086465 |
| 0 | 0.2249730252 | 0 | 0.2168180342 | 0.5582089406 |
| 0 | 0 | 0.4904068955 | 0 | 0.5095931045 |
| 0.2310296614 | 0 | 0.5442016106 | 0.224768728 | 0 |
| 0.01739861546 | 0.2088879599 | 0.2223130609 | 0.5471158311 | 0.004284532612 |
| 0 | 0.5309515958 | 0.2319648264 | 0.2370835777 | 0 |
| 0.4959614665 | 0 | 0 | 0.5040385335 | 0 |
| 0.5363785252 | 0.227062008 | 0 | 0.2365594669 | 0 |
| 0 | 0 | 0.7809282607 | 0.2190717393 | 0 |
| 0.2161803028 | 0.2176035052 | 0.02451466236 | 0.5417015297 | 0 |
| 0.1253980405 | 0.1151854478 | 0.1282873892 | 0.1098044354 | 0.5213246871 |
| 0.2206686281 | 0 | 0.5490565201 | 0.00515155436 | 0.2251232974 |
| 0.2505882319 | 0.2593855362 | 0 | 0.2478813235 | 0.2421449084 |
| 0.5476407458 | 0 | 0.221599459 | 0 | 0.2307597952 |
| 0.2164014906 | 0.2311595894 | 0 | 0.001391654508 | 0.5510472655 |
| 0 | 0 | 0 | 0 | 0 |

**Supplementary Table 4: Antibodies**

| Target | Colour | Supplier | Reference | Dilution | Application |
|---|---|---|---|---|---|
| f-actin | Atto550 | Sigma-Aldrich | 19083 | 1:300 | IF |
| desmin (rabbit) | - | Abcam | ab227651 | 1:400 | IF |
| myosin | - | Abcam | ab51263 | 1:2,000 | IF |
| myosin | - | Abcam | ab11083 | 1:5,000 | Jess |
| mouse | AF488 | Invitrogen | A-11001 | 1:1,000 | IF |
| rabbit | DyLight550 | Thermo Fisher Scientific | SA5-10033 | 1:250 | IF |
